# Type I interferon-dependent CCL4 is induced by a cGAS/STING pathway that bypasses viral inhibition and protects infected tissue, independent of viral burden

**DOI:** 10.1101/616888

**Authors:** Nikhil J. Parekh, Tracy E. Krouse, Irene E. Reider, Ryan P. Hobbs, Brian M. Ward, Christopher C. Norbury

**Affiliations:** Department of Microbiology and Immunology, The Pennsylvania State University College of Medicine, Hershey, PA 17033, USA; Department of Dermatology, The Pennsylvania State University College of Medicine, Hershey, PA 17033, USA; Department of Microbiology and Immunology, University of Rochester Medical Center, Rochester, NY 14642, USA

## Abstract

Type I interferons (T1-IFN) are critical in the innate immune response, acting upon infected and uninfected cells to initiate an antiviral state by expressing genes that inhibit multiple stages of the lifecycle of many viruses. T1-IFN triggers the production of Interferon-Stimulated Genes (ISGs), activating an antiviral program that reduces virus replication. The importance of the T1-IFN response is highlighted by the evolution of viral evasion strategies to inhibit the production or action of T1-IFN in virus-infected cells. T1-IFN is produced via activation of pathogen sensors within infected cells, a process that is targeted by virus-encoded immunomodulatory molecules. This is probably best exemplified by the prototypic poxvirus, Vaccinia virus (VACV), which uses at least 6 different mechanisms to completely block the production of T1-IFN within infected cells *in vitro*. Yet, mice lacking aspects of T1-IFN signaling are often more susceptible to infection with many viruses, including VACV, than wild-type mice. How can these opposing findings be rationalized? The cytosolic DNA sensor cGAS has been implicated in immunity to VACV, but has yet to be linked to the production of T1-IFN in response to VACV infection. Indeed, there are two VACV-encoded proteins that effectively prevent cGAS-mediated activation of T1-IFN. We find that the majority of VACV-infected cells *in* vivo do not produce T1-IFN, but that a small subset of VACV-infected cells *in vivo* utilize cGAS to sense VACV and produce T1-IFN to protect infected mice. The protective effect of T1-IFN is not mediated via ISG-mediated control of virus replication. Rather, T1-IFN drives expression of CCL4, which recruits inflammatory monocytes that constrain the VACV lesion in a virus replication-independent manner by limiting spread within the tissue. Our findings have broad implications in our understanding of pathogen detection and viral evasion *in vivo*, and highlight a novel immune strategy to protect infected tissue.

**Summary:** The recognition of virus infection leads to a quick and robust antiviral response mediated by type I interferons (T1-IFN). Nearly all viruses have acquired genes that block the induction or action of T1-IFN in order to attain a replicative advantage. Some viruses thwart the T1-IFN response so thoroughly, that cells infected *in vitro* do not produce any T1-IFN. And yet, animal models with defects in T1-IFN signaling are more sensitive to infection with these viruses than their wild-type counterparts. In this study, we find evidence to explain these otherwise contradicting findings. We show that a small population of infected cells *in vivo* are able to utilize a pathogen-sensing pathway that is completely blocked *in vitro*. T1-IFN produced by these specialized cells protects mice by recruiting inflammatory monocytes that restrict the spread of virus within infected tissue, independent of viral burden. Our findings have a direct impact on our understanding of how viruses are detected in infected tissue, and present a novel strategy of the immune system to limit pathology at peripheral sites of infection.

## Introduction

Type I interferons (T1-IFN) are a critical component of the response to many viruses [1]. First described in the 1950’s, cells infected with a virus produce T1-IFNs which induce an antiviral state in surrounding cells [2, 3]. All T1-IFNs signal through a common receptor, the T1-IFN receptor (IFNαR), and consist of a single IFN-β, about a dozen similar IFN-α, and IFN-ε, κ, ζ, δ, τ, ω, and ν [4]. T1-IFN production is triggered by recognition of viral proteins or nucleic acids by host Pattern Recognition Receptors (PRRs) [5]. PRRs can be cell surface or endosomal Toll-like receptors (TLRs) or cytosolic nucleic acid sensors. IFNαR signaling initiates a positive feedback loop that produces more T1-IFN and expression of the full breadth of IFN-α subtypes [6]. The antiviral state is the effect of hundreds of Interferon-Stimulated Genes (ISGs) driven by IFNαR signaling [7, 8], which function to inhibit virus entry, cell growth, and virus replication. This is the major mechanism that controls virus replication and host tissue pathology early after infection.

The importance of the T1-IFN pathway in antiviral defense is underscored by two observations. First, animals lacking IFNαR are highly susceptible to virus infection [1, 9]. Second, the majority of animal viruses encode at least one, but usually many, immunomodulatory proteins that interfere with, and often completely block, the production or action of T1-IFN [10, 11]. If viruses have evolved to block T1-IFN so efficiently, why is T1-IFN so important in resistance to virus infection *in vivo*? Here, we compare the *in vitro* and *in vivo* responses to a virus that infects peripherally to answer this question.

Viruses have evolved to evade the T1-IFN response using multiple strategies. First, many viruses can block the pathways that sense infection and trigger T1-IFN production by preventing PRRs from accessing their ligands, or by blocking PRR activation or signaling [10, 11]. Viruses can also produce soluble cytokine-binding proteins that bind to, and prevent the action of, secreted T1-IFN upon neighboring cells, and can also encode proteins that inhibit signaling through IFNαR within infected cells, preventing ISG expression [11, 12]. However, despite all of these approaches deployed by viruses to subvert the host interferon response, the T1-IFN system remains an important effector arm of the innate immune response following virus infection.

We have utilized a viral infection model which exemplifies the ability of viruses to evade the T1-IFN response, the prototypic poxvirus vector, Vaccinia virus (VACV). We find a complete blockade in production of T1-IFN by VACV-infected cells *in vitro* [13, 14]. VACV has been reported to block the action of multiple cell surface, endosomal or cytosolic PRRs, including the predominant cytosolic DNA sensor cyclic GMP-AMP synthase (cGAS) and downstream signal transduction by its adapter protein Stimulator of Interferon Genes (STING) [15-18]. Indeed, we find evidence of efficient action of multiple mechanisms of viral evasion *in vivo*. Nevertheless, T1-IFN is still produced by a subset of infected cells *in vivo* in a cGAS- and STING-dependent manner. The T1-IFN produced protects mice against tissue pathology caused by VACV infection. However, the protective effect of T1-IFN is not mediated by the classical mechanism of control of virus replication, as neither ISG induction nor VACV replication are altered in the absence of T1-IFN signaling. Rather, it is mediated through T1-IFN-dependent production of the chemokine CCL4, which is required for localization of inflammatory monocytes to the VACV lesion. Monocytes enter the VACV lesion, where they become non-productively infected, allowing them to soak up excess virions and prevent spread to neighboring cells. Therefore, in spite of all the mechanisms deployed against T1-IFN by viruses, the host stays ahead of pathogen evolution by deploying alternative strategies to both produce T1-IFN, and to mediate protective strategies downstream of T1-IFN signaling.

## Results

### T1-IFN is not expressed within infected cells *in vitro*, but T1-IFN expression is required *in vivo*

The primary cell types infected with VACV after immunization in the skin are keratinocytes and recruited monocyte/macrophages [19]. We infected representative cell lines with VACV and measured the production of IFN-β. Neither a keratinocyte line (**Fig. 1A**) nor a macrophage cell line (**Fig. 1B**) produced T1-IFN, measured by mRNA or protein, following exposure to live VACV. However, both cell lines produced significant quantities of IFN-β mRNA and protein following exposure to controls, namely heat-inactivated VACV or Vesicular stomatitis virus (**Fig. 1A, B**). Therefore, the expression of multiple (≥ 6) immune evasion proteins blocked T1-IFN production from cells present at the site of infection in the skin in response to live VACV [15, 16, 20].

**Figure 1.**
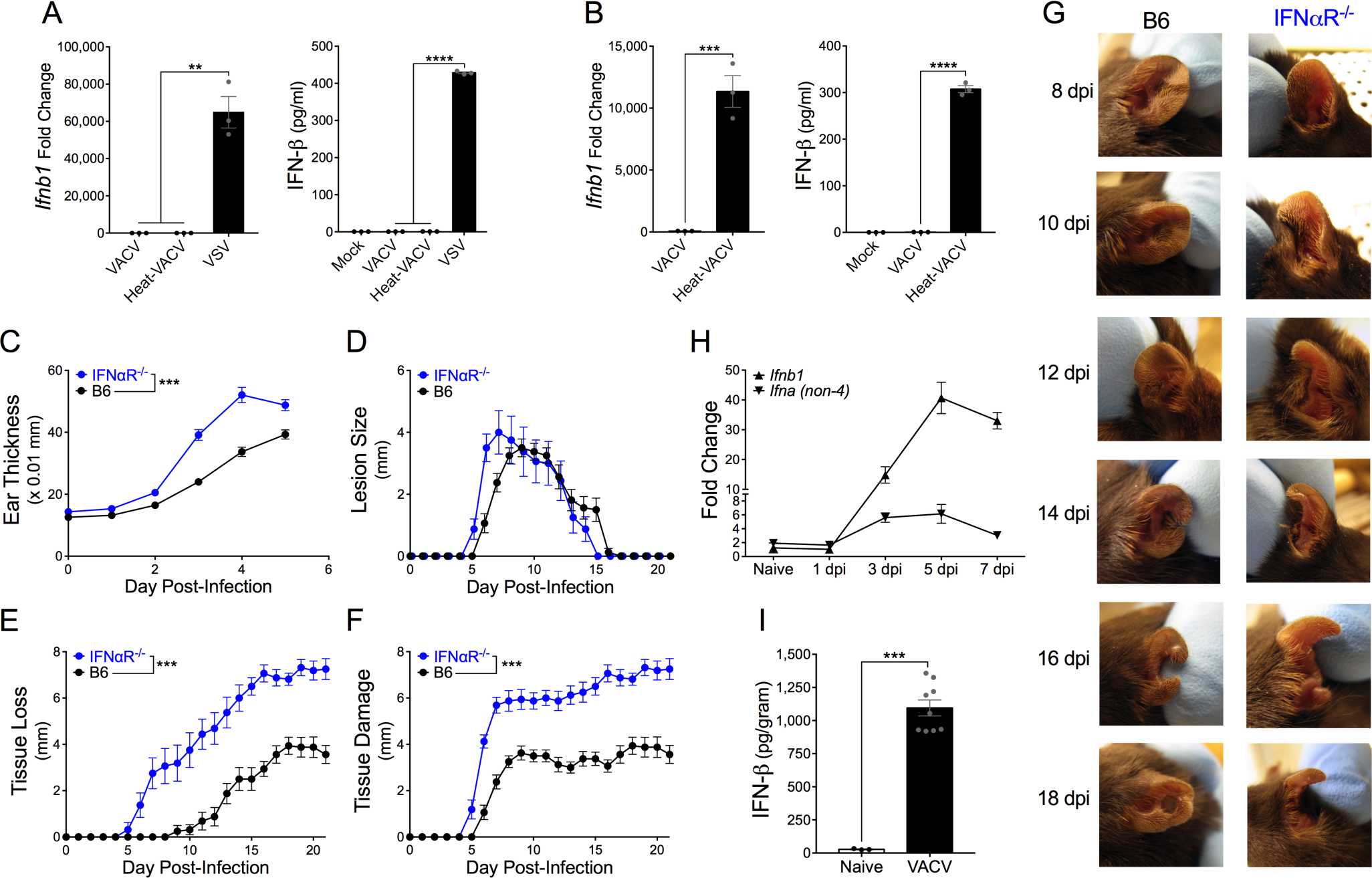
T1-IFN is not expressed within infected cells *in vitro*, but T1-IFN expression is required *in vivo*. (A,B) Expression of IFN-β from murine a keratinocyte (A) or macrophage (B) cell line measured by RT-qPCR 6 hpi (left) or ELISA 24 hpi (right). (C-F) Tissue pathology of the ear pinnae after intradermal infection with VACV; ear thickness (C), lesion size (D), tissue loss (E), or total tissue damage (F), n=8 mice per group. (G) Representative pictures of ear pinnae following intradermal VACV infection. (H,I) Expression of T1-IFN in ear tissue of wild-type mice at the indicated time post-infection measured by RT-qPCR (H, n=3 naïve, 4 VACV-infected mice per group), or 5 dpi by ELISA (I).

In marked contrast, when we infected WT mice or mice lacking the T1-IFN receptor (IFNαR^-/-^) with VACV intradermally in the ear [21, 22] there was a significant difference in the tissue response to the virus. This was apparent as early as 3 days post-infection, when swelling at the infection site was greater in IFNαR^-/-^ mice (**Fig. 1C**), and progressed to produce a larger VACV lesion (**Fig. 1D**), greater tissue loss (**Fig. 1E**) and overall tissue damage (lesion size + tissue loss) (**Fig. 1F**) in IFNαR^-/-^ mice. **Figure 1G** shows that VACV infection causes only a small hole in the tissue of wild-type mice, but the majority of the ear is lost following infection of IFNαR^-/-^ mice. Therefore, T1-IFN signaling is clearly required to moderate tissue damage following dermal VACV infection *in vivo*. To measure T1-IFN production *in vivo*, we harvested tissue from VACV-infected ears and assayed for induction of IFN-β and IFN-α mRNA. Even when utilizing stringent RNA purification procedures, no significant induction of IFN mRNA could be detected prior to 3 days post-infection (**Fig. 1H**). However, expression of IFN-β and IFN-α mRNA could be detected at 3 days post-infection and peaked at 5 days post-infection (**Fig. 1H**). We also detected IFN-β protein by ELISA from ear tissue at the peak of mRNA production, 5 days after dermal infection (**Fig. 1I**). Therefore, despite the blockade of T1-IFN production in infected cells by immune modulators *in vitro*, VACV induces T1-IFN *in vivo*, and IFNαR signaling is required to prevent excessive tissue pathology following VACV infection.

### A positive feedback loop is required for T1-IFN production in response to VACV

IFN-β and the different IFN-α subtypes have differential effects on signals transduced through the IFNαR [23]. To understand the profile of T1-IFN genes induced by VACV infection, we assayed the expression of representative genes within the T1-IFN family in the ear 5 days post-infection with VACV. IFN-β was strongly induced in all experiments (**Fig. 2A**) but, although IFN-α2 is the predominant gene present in many arrays and multiplex assays, we found that it was not induced by VACV (**Fig. 2A**). Rather, IFN-α4 and, to a lesser but modest extent, IFN-α5 were the only IFN-α species induced by dermal VACV infection (**Fig. 2A**). In most cells, robust T1-IFN production requires positive feedback signaling through IFNαR [24, 25]. We examined induction of T1-IFN in mice lacking IFNαR signaling and found poor induction of both IFN-β and IFN-α4 mRNA in these mice, compared to wild-type controls (**Fig. 2B**). IFNαR stimulation results in phosphorylation of the transcription factors Signal Transducer and Activator of Transcription (STAT) STAT1 and STAT2, which form a trimeric complex with IFN Regulatory Factor (IRF) IRF9, and are required to transduce T1-IFN signals [26, 27]. Both STAT1 (**Fig. 2C**) and STAT2 (**Fig. 2D**) were required for optimal induction of T1-IFN. Therefore, although virus-encoded proteins block T1-IFN from binding its receptor and the downstream activation and function of STAT proteins, a positive feedback loop is required for an efficient T1-IFN response following dermal VACV infection.

**Figure 2.**
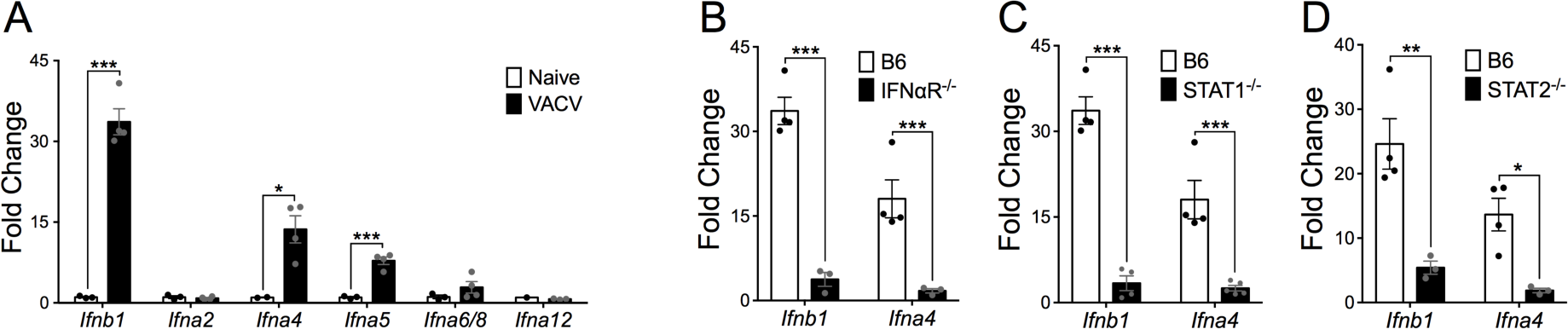
A positive feedback loop is required for T1-IFN production in response to VACV. (A) Expression of T1-IFN subtypes in ear tissue of wild-type mice 5 dpi measured by RT-qPCR. (B-D) Expression of early T1-IFNs in ear tissue of mice 5 dpi measured by RT-qPCR.

### cGAS sensing of VACV-infected cells induces T1-IFN to limit local tissue pathology

Induction of T1-IFN requires recognition of viral components by PRRs and signal transduction through adapter proteins. Cell surface and endosomal TLRs utilize either Myeloid Differentiation Primary Response 88 (MyD88) or TIR-domain-containing adapter-inducing interferon-β (TRIF) proteins to induce T1-IFN. Cytosolic nucleic acid sensors utilize the Mitochondrial Antiviral Signaling protein (MAVS) or STING for signal transduction. Each of these pathways is regularly targeted by a host of viruses, including poxviruses, which alter recognition by, or signaling downstream from, the MyD88 [28], TRIF [29], MAVS [20, 30] or STING [15, 16] pathways. We have previously found that MyD88 is required for optimal immune responses to dermal VACV infection [31], and others have found that MyD88 signaling is crucial in NK [32], T cell [33, 34] and Plasmacytoid DC [35] responses. However, we found that MyD88 was not required for the induction of IFN-β in response to VACV (**Fig. 3A**). Although TRIF has been implicated in limiting VACV replication within infected macrophages [36], we found no defect in the induction of IFN-β in response to VACV in mice with a non-functional TRIF gene (**Fig. 3B**). Deletion of the virus-encoded immunomodulatory protein E3L can rescue IFN production *in vitro* via recognition of dsRNA in a manner dependent on the PRR-adapter protein MAVS [20]. However, *in vivo*, MAVS-deficient mice were also able to effectively induce IFN-β in response to VACV (**Fig. 3C**). VACV is a large dsDNA virus that carries out its lifecycle entirely in the cytoplasm of infected cells, and thus is likely sensed by cytosolic DNA sensors upstream of the adapter protein STING. Even though VACV efficiently blocks sensing via the STING pathway *in vitro* [15-17], induction of IFN-β was heavily reliant on STING signaling (**Fig. 3D**), indicating that recognition of VACV DNA was likely responsible for T1-IFN production.

**Figure 3.**
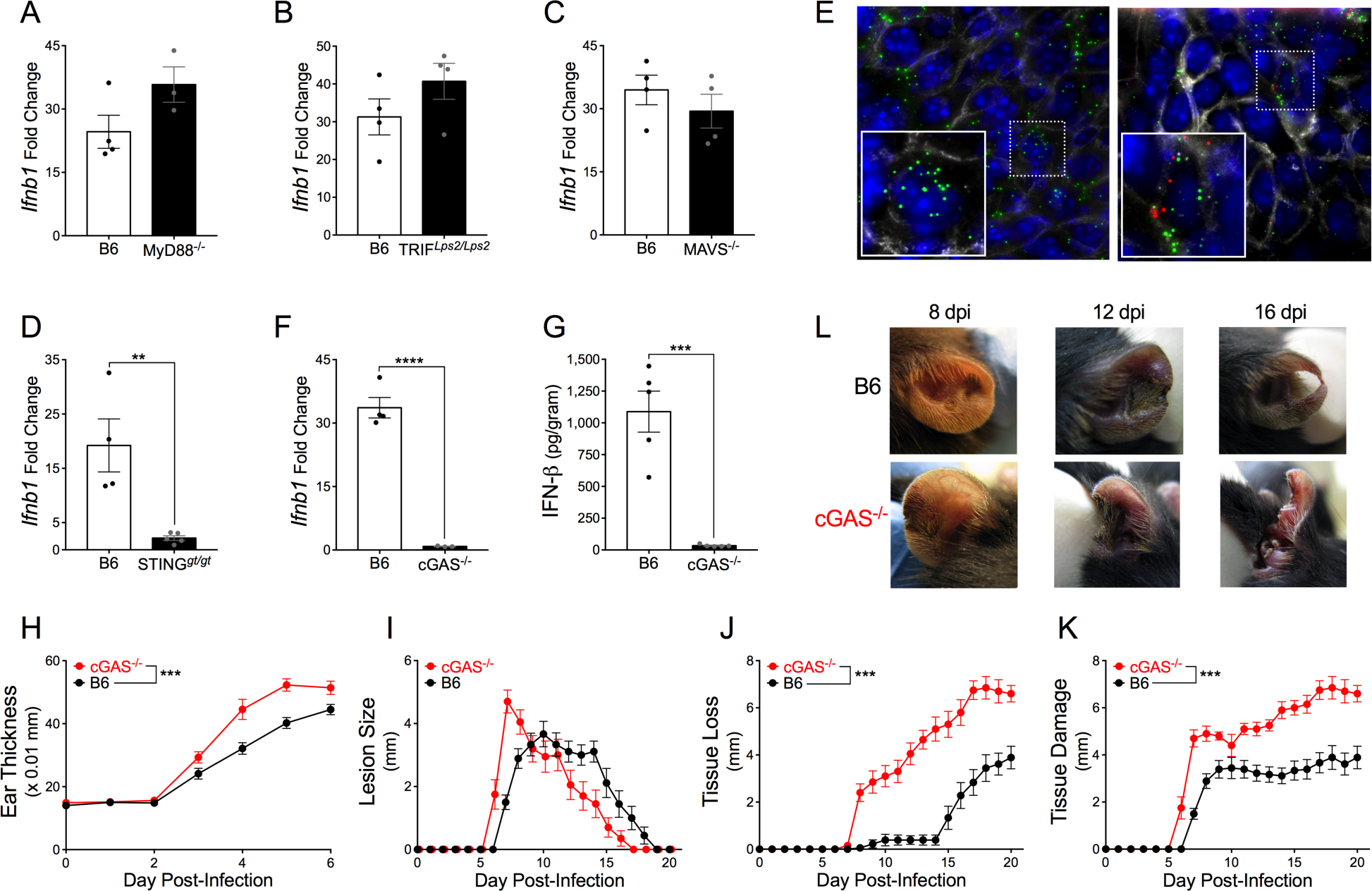
cGAS sensing of VACV-infected cells induces T1-IFN to limit local tissue pathology. (A-D) Expression of IFN-β in ear tissue of mice 5 dpi measured by RT-qPCR. (E) Cryosections of ear tissue from wild-type mice 5 dpi with VACV-OVA stained with Wheat-Germ Agglutinin (Grey) to visualize cell membranes and DAPI (Blue) for nuclei. RNA fluorescence *in situ* hybridization was used to detect viral RNA (OVA-Green) and cellular IFN-β (Red). (F,G) Expression of IFN-β in ear tissue of mice 5 dpi measured by RT-qPCR (F) or ELISA (G). (H-K) Tissue pathology of the ear pinnae after intradermal infection with VACV; ear thickness (H), lesion size (I), tissue loss (J), or total tissue damage (K), n=9-10 mice per group. (L) Representative pictures of ear pinnae following intradermal VACV infection.

The almost total requirement for STING signaling was curious, as cells infected with VACV *in vitro* do not produce T1-IFN (**Fig. 1**). Therefore, we considered the possibility that T1-IFN may only be produced by small numbers of uninfected cells, which are not subject to the same strict immunomodulation as VACV-infected cells. We infected mice with a recombinant VACV expressing chicken egg ovalbumin (OVA), and on day 5 post-infection conducted *in situ* hybridization with probes targeting either OVA (to identify VACV-infected cells) or IFN-β mRNA. To allow accurate definition of cell boundaries, we stained with fluorescently-labeled wheat-germ agglutinin, which efficiently labeled all cell-surface membranes in the infected ear. In the majority of areas in which we could observe OVA mRNA there was little or no IFN-β mRNA visible (**Fig. 3E, left panel**), raising the possibility that it was indeed uninfected cells that were the source of IFN-β. However, in multiple experiments we never observed IFN-β mRNA in cells that did not also display OVA mRNA (**Fig. 3E, right panel**). Rather, it was a small, spatially distinct subpopulation of VACV-infected, OVA mRNA-expressing cells that produced IFN-β mRNA within the cytosol (**Fig. 3E**). Therefore, the inhibitory effect of VACV upon STING signaling that appears complete *in vitro*, is bypassed *in vivo*, allowing T1-IFN production.

Upstream of STING, cGAS has been implicated in the recognition of multiple virus infections, including VACV [18, 37, 38]. However, in addition to its well established role in induction of T1-IFN production, cGAS can also activate NF-*κ*B [39-41] and STAT6 [42, 43], both of which can impact the anti-poxvirus immune response [44-49]. We found that induction of IFN-β mRNA and protein was impaired in cGAS-deficient mice to a similar extent to that observed in mice that lacked STING signaling (**Fig. 3F, G**). To investigate a role for cGAS-induced T1-IFN during dermal VACV infection, we measured local tissue pathology in cGAS^-/-^ mice compared to wild-type mice after intradermal infection of the ear. cGAS^-/-^ mice displayed increased tissue thickness beginning by 3 days post-infection (**Fig. 3H**), the time point at which we first measured induction of T1-IFN transcripts (**Fig. 1H**). Similar to the phenotype observed in IFNαR^-/-^ mice, cGAS^-/-^ mice also exhibited larger lesions, greater tissue loss and total tissue damage compared to wild-type mice (**Fig. 3I-L**). Therefore, cGAS-mediated production of T1-IFN is predominantly responsible for moderating tissue pathology following dermal VACV infection. Thus, although cGAS-mediated induction of T1-IFN is completely blocked *in vitro*, the host uses this pathway to induce tissue protective T1-IFN *in vivo*, bypassing virus-mediated inhibitory pathways.

### T1-IFN does not control VACV replication in the skin

Many ISGs have direct antiviral properties that can blunt propagation of virus within infected tissue [7]. Therefore, it was important to examine whether T1-IFN-mediated upregulation of antiviral ISGs in the ear was responsible for control of VACV replication, and thus whether a lack of ISG induction was responsible for the tissue pathology we observed after infection of IFNαR^-/-^ mice. We examined the transcript levels of 81 prominent antiviral ISGs 5 days after dermal VACV infection by RT-qPCR array and found strong (>4-fold to ∼300-fold) induction of a number of ISGs (**Fig. 4A**). Surprisingly, we found that a small number of IFN-responsive transcripts were downregulated upon dermal VACV infection, including *irf4, irf5* and *ifi27* (**Fig. 4A**). We also found robust induction of a number of antiviral ISGs including *Ifit1, Ifi204, ISG15, OAS1, Ifih1, irf1, irf7* and *irf8* (**Fig. 4A**). However, we did not find that induction of any of the antiviral ISGs was dependent upon T1-IFN signaling (**Fig. 4B**). This suggests that redundant pathway(s) exists for induction of these antiviral genes. In addition, it clearly demonstrates that upregulation or downregulation of these antiviral ISGs by T1-IFN signaling is not responsible for the tissue pathology phenotype we observed in IFNαR^-/-^ and cGAS^-/-^ mice.

**Figure 4.**
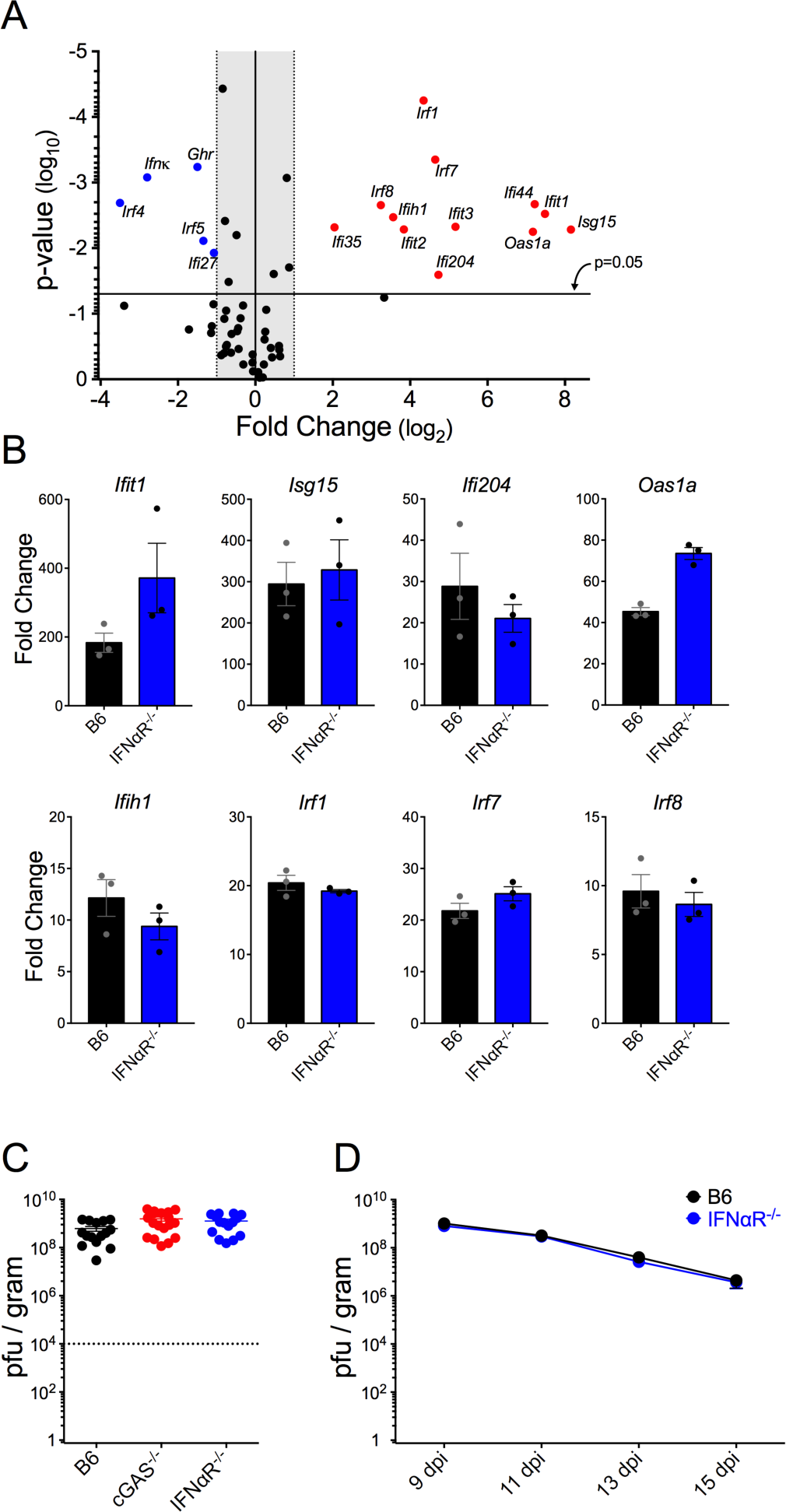
T1-IFN does not control VACV replication in the skin. (A) Expression of antiviral ISGs in ear tissue of wild-type mice 5 dpi compared to naïve controls measured using RT-qPCR array. (B) Expression of induced ISGs from (A) in ear tissue of wild-type and IFNαR^-/-^ mice 5 dpi measured using RT-qPCR array. (C) VACV titer in ears of mice 5 days post-intradermal infection with VACV. Dotted line indicates initial inoculum. (D) VACV titer in ears of mice at the indicated day post-infection with VACV by scarification, n=5 mice per group.

To directly address the role for T1-IFN in control of VACV replication *in vivo*, we compared VACV titer in the ears of cGAS^-/-^ and IFNαR^-/-^ mice to wild-type mice at 5 days post-infection, the peak of VACV replication [50]. Both IFNαR^-/-^ and cGAS^-/-^ mice displayed only a modest 2-fold increase in infectious VACV titer (**Fig. 4C**), which is unlikely to explain the drastic change in pathology. VACV gene expression and infectious titer decrease after 5 days post-infection [19, 50], so it is possible that T1-IFN controls viral clearance rather than limiting peak VACV titers. However, the tissue loss exhibited by IFNαR^-/-^ mice makes it difficult to accurately evaluate VACV titer at late times post-intradermal infection. In order to assess the role of T1-IFN in clearance of VACV, we infected the ears of IFNαR^-/-^ and wild-type mice with VACV by scarification, an infectious methodology that results in a milder pathology. We found that the kinetics of VACV clearance was unaffected in the absence of T1-IFN signaling (**Fig. 4D**), so the protective effect of T1-IFN during dermal VACV infection is independent of its direct antiviral characteristic.

### T1-IFN induces *ccl4* to recruit protective monocytes to the site of infection

In addition to the induction of antiviral ISGs, it is now becoming clear that T1-IFN can modulate the innate and adaptive immune response, independent of any effects upon virus replication [51, 52]. Therefore, we used a second qPCR array analysis to compare the change in expression of 84 cytokines and chemokines in the ear 5 days after dermal VACV infection. We found strong infection-mediated induction of 28 of 84 transcripts, with some that are known to be interferon-responsive, including the CXCR3 ligands *cxcl9* and *cxcl10*, as well as IFN-γ (**Fig. 5A**). To investigate which of these transcripts were dependent upon T1-IFN signaling, we harvested mRNA from ear tissue of VACV-infected wild-type and IFNαR^-/-^ mice and performed a similar qPCR array analysis. Although VACV strongly induced many transcripts in wild-type mice, we found that the expression of only one transcript was differentially modulated >2-fold in a statistically significant manner in the absence of T1-IFN signaling. The IFNαR-dependent gene was *ccl4* (macrophage inflammatory protein-1β) (**Fig. 5B**). We were able to confirm that expression of *ccl4* was dependent on both IFNαR signaling and cGAS expression using a targeted qPCR assay (**Fig. 5C**).

**Figure 5.**
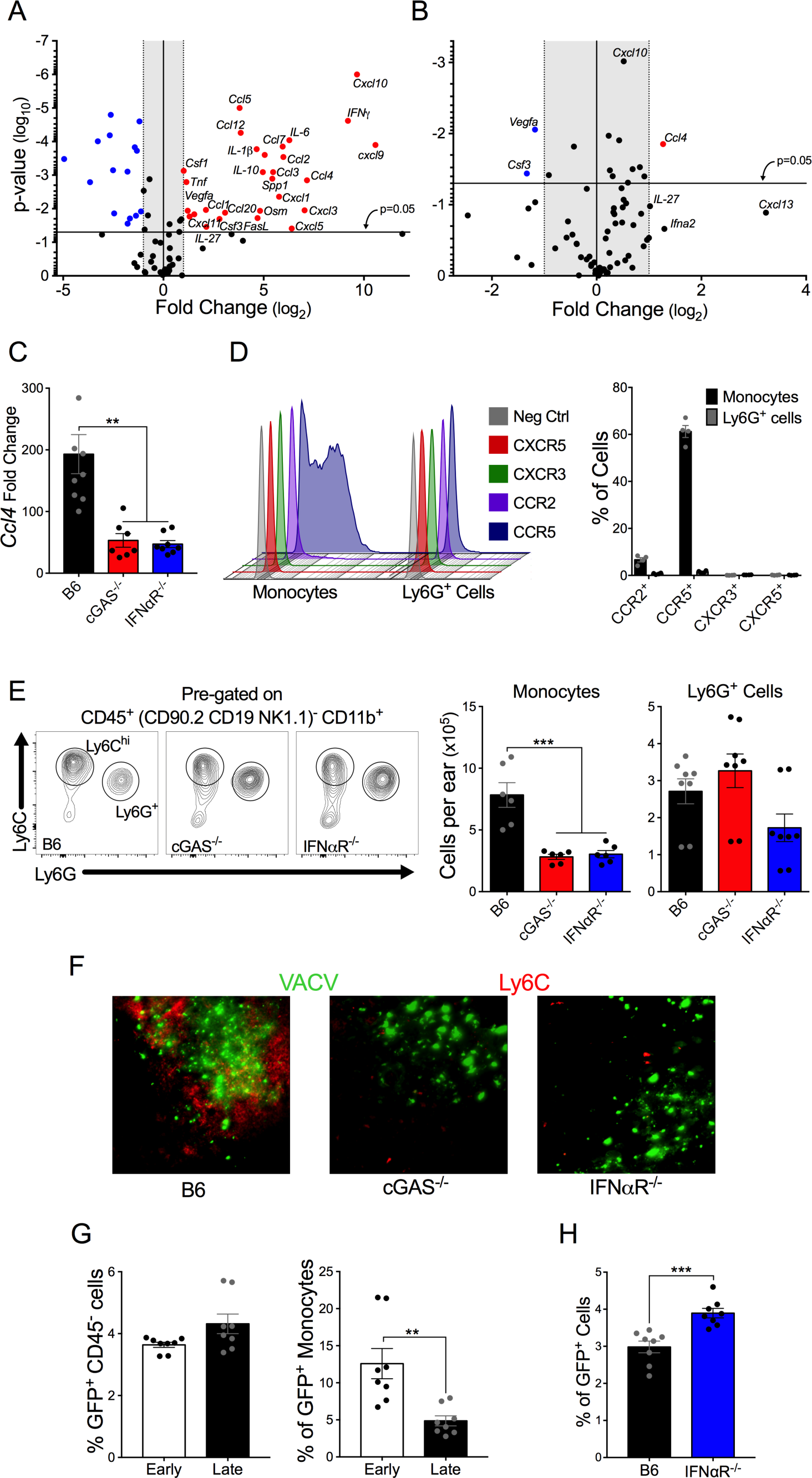
T1-IFN induces *ccl4* to recruit protective monocytes to the site of infection. (A,B) Expression of cytokines and chemokines in ear tissue of mice 5 dpi measured using RT-qPCR array. Gene expression changes in wild-type mice in response to VACV (A), or gene expression in VACV-infected wild-type mice compared to VACV-infected IFNαR^-/-^ mice (B), n=3 naïve, 9 VACV-infected mice per group. (C) Expression of *ccl4* in ear tissue of VACV-infected mice 5 dpi measured by RT-qPCR. (D) Chemokine receptor expression on inflammatory monocytes and Ly6G^+^ cells recruited into VACV-infected ears 5 dpi. Representative flow cytometry plots (left) and proportion of cells expressing each receptor (right). (E) Myeloid cell populations in VACV-infected ears 5 dpi measured by flow cytometry. Representative flow cytometry plots (left) and absolute number of cells (right). (F) Immunofluorescence microscopy of ears 5 dpi with VACV-NP-S-eGFP reveal monocyte localization (Ly6C-Red) in relation to VACV-infected cells (GFP-Green). (G) Percentage of GFP^+^ cells in the ear 5dpi with VACV Early-GFP or VACV Late-GFP measured by flow cytometry. (H) Percentage of GFP^+^ cells in the ear 5dpi with VACV-NP-S-eGFP measured by flow cytometry.

The cellular response to dermal VACV infection is detected at 3 days post-infection when inflammatory monocytes move into the ear, and numbers of these cells peak at 5 days post-infection [50], i.e. with similar kinetics to T1-IFN induction (**Fig. 1H**). Recruitment of classical inflammatory monocytes occurs with similar kinetics to the induction of T1-IFN transcripts, and is followed by recruitment of a non-classical tissue-protective Ly6G^+^ cell population that primarily occurs after the peak of T1-IFN production [50]. Both classical inflammatory monocytes and the non-classical tissue-protective Ly6G^+^ cell population home to foci of VACV infection, although only classical inflammatory monocytes are infected within the lesion [19]. Local depletion of either inflammatory monocytes [53] or the tissue-protective Ly6G^+^ cells [50] produces tissue pathology similar to that observed in cGAS^-/-^ or IFNαR^-/-^ mice. However, depletion of T cells does not affect local pathology or systemic spread of virus following dermal VACV infection [50, 53]. Because the recruitment of classical inflammatory monocytes occurs concurrently with the induction of T1-IFN gene transcripts (**Fig. 1H**), and because the onset of cell recruitment into the ear coincides with the onset of increased tissue pathology in mice lacking T1-IFN signaling (**Fig. 1C-F**), we sought to determine if T1-IFN-induced *ccl4* plays a role in the recruitment of classical inflammatory monocytes, as well as the later recruitment of the Ly6G^+^ cell population.

We assayed for the presence of chemokine receptors on the cell surface of inflammatory monocytes and tissue-protective Ly6G^+^ cells that traffic into VACV-infected ears. From our array data, we found a statistically significant change in the expression of *ccl4*, a large, but non-statistically significant change in expression of *cxcl13* as well as a slight, significant change in the expression of *cxcl10* (**Fig. 5B**). Using flow cytometry, we assayed for cell-surface expression of the receptors for those respective chemokines, as well as for the prototypic monocyte chemokine receptor, CCR2, the receptor for CCL2 that was upregulated upon VACV infection (**Fig. 5A**). We found that the majority of CD11b^+^ LyC^hi^ Ly6G^-^ CD64^+^ inflammatory monocytes present in the ear of VACV-infected mice at 5 days post-infection were CCR5^+^, the receptor for CCL4, rather than CCR2^+^, or positive for the receptors for CXCL13 (CXCR5) or CXCL9 and CXCL10 (CXCR3) (**Fig. 5D**).

We next examined whether the deficiency in *ccl4* expression by the absence of T1-IFN prevented recruitment of CCR5^+^ inflammatory monocytes to the VACV-infected ear. We assayed myeloid cell infiltration in cGAS^-/-^ and IFNαR^-/-^ mice and found a defect in infiltration of inflammatory monocytes, but not Ly6G^+^ cells, into the infected ear compared to wild-type mice (**Fig. 5E**). A small number of inflammatory monocytes still moved into the infected ear, however, Ly6C^+^ inflammatory monocytes failed to localize to the VACV lesion in the absence of T1-IFN induction or signaling (**Fig. 5F**). Therefore, cGAS-mediated induction of T1-IFN during dermal VACV infection is required for *ccl4* expression and CCR5-dependent recruitment of inflammatory monocytes to viral lesions.

Recruited inflammatory monocytes moderate pathology following VACV infection in a manner independent of control of virus replication and expression of antiviral ISGs. However, the mechanisms used by monocytes/macrophages to mediate protective antiviral responses remain poorly described at present. Following infection with some viruses, macrophage populations are preferentially infected in order to facilitate initiation of an adaptive immune response [54-56] or otherwise restrict virus spread [53, 57, 58]. However, primary macrophage populations do not typically support VACV replication, restricting gene expression at a point prior to DNA replication [59]. Therefore, we sought to explore whether infected inflammatory monocytes restricted VACV replication *in vivo* using a similar mechanism. We infected wild-type mice with VACV that expresses GFP driven by either the early/late promoter p7.5 (Early-GFP) or the late promoter p11 (Late-GFP), which is active only after DNA replication, and assayed GFP expression in inflammatory monocytes in the ear 5 days post-infection. In CD45^-^ cell populations there were equivalent numbers of cells expressing early and late promoter-driven GFP, indicating that there was no blockade in VACV replication within these cells (**Fig. 5G, left panel**). However, a greater proportion of inflammatory monocytes expressed early promoter-driven GFP than GFP driven by the late p11 promoter (**Fig. 5G, right panel**), indicative of a blockade of VACV replication in the inflammatory monocyte population. Therefore, inflammatory monocytes within the VACV lesion can potentially act as a “virus sponge,” becoming infected to spare nearby cells from infection. To examine whether this occurred, we infected wild-type or IFNαR^-/-^ mice with VACV that expresses GFP that accumulates in the nucleus of infected cells (VACV-NP-S-eGFP). IFNαR^-/-^ mice had a greater proportion of VACV-infected cells in the ear 5 days post-infection compared to wild-type mice (**Fig. 5H**). Thus, inflammatory monocytes enter the VACV lesion, become non-productively infected, and prevent the infection of neighboring cells. These data support a role for T1-IFN-mediated restriction of virus spread and subsequent tissue damage via a novel immunomodulatory mechanism that does not require a direct effect upon virus replication via the expression of antiviral ISGs.

### *ccl4* expression restores inflammatory monocyte recruitment and ameliorates tissue damage

To examine the importance of T1-IFN-mediated *ccl4* expression in the control of VACV-mediated tissue damage in the dermis, we constructed a recombinant VACV that expresses a construct driven by the relatively weak C6 promoter containing murine ccl4 and the red fluorescent protein mKate2 (VACV-CCL4) (**Fig. 6A**). Upon infection of cells *in vitro*, the rVACV produced mKate (**Fig. 6B**), as well as *ccl4* mRNA (**Fig. 6C**) and protein (**Fig. 6D**). To examine whether VACV-driven expression of *ccl4* could rescue recruitment of inflammatory monocytes, we infected IFNαR^-/-^ mice with VACV-CCL4 or a control VACV that did not express *ccl4*. Five days after infection we assayed the recruitment of CD11b^+^ LyC^hi^ Ly6G^-^ CD64^+^ inflammatory monocytes or tissue protective Ly6G^+^ myeloid cells in the infected ear. VACV-expression of CCL4 increased the number and proportion of inflammatory monocytes in the ears of VACV-infected mice in the absence of T1-IFN signaling (**Fig. 6E**) without altering recruitment of tissue-protective Ly6G^+^ myeloid cells (**Fig. 6F**). Further, and more importantly, restoration of *ccl4* expression was able to rescue localization of Ly6C^+^ inflammatory monocytes to foci of virus infection in IFNαR^-/-^ mice (**Fig. 6G**). To examine whether CCL4-mediated inflammatory monocyte recruitment ameliorated the tissue damage we observed in mice lacking IFNαR signaling, we infected IFNαR^-/-^ mice with VACV-CCL4 or a control VACV that did not express *ccl4* and monitored multiple aspects of tissue damage as outlined above. We found a statistically significant reduction in ear swelling, lesion size and the combined criteria of total tissue damage (**Fig. 6H-K**). These data strongly support a requisite role for CCL4-mediated recruitment of inflammatory monocytes in moderating tissue damage, but not virus replication, following dermal VACV infection.

**Figure 6.**
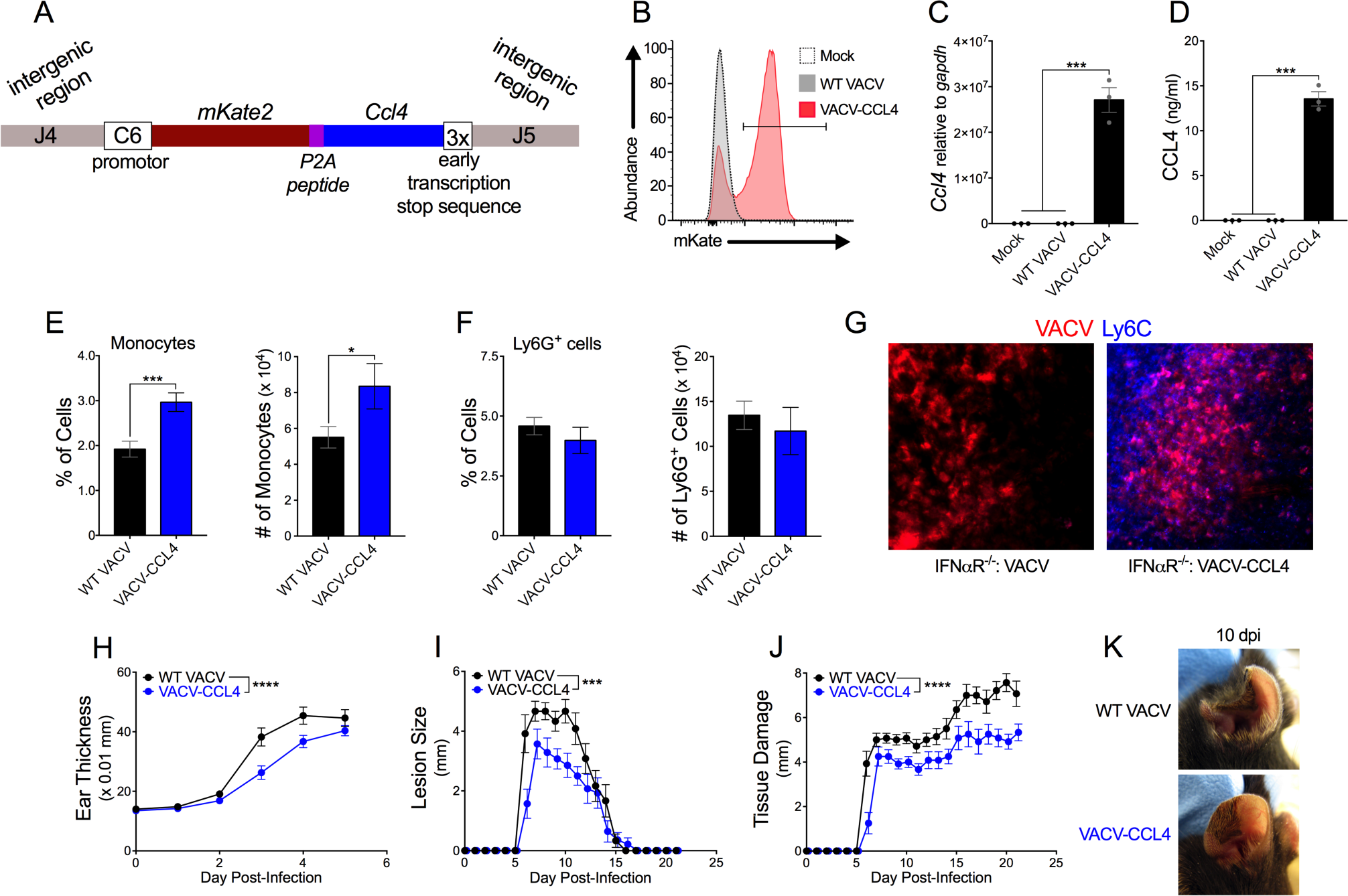
*ccl4* expression restores inflammatory monocyte recruitment and ameliorates tissue damage. (A) Schematic of rVACV expressing *ccl4* and mKate2. (B) Detection of mKate2 expression in 143B cells 6 hpi assayed using flow cytometry. (C) Expression of murine *ccl4* in 143B cells 6 hpi measured by RT-qPCR. (D) Secretion of murine CCL4 from 143B cells 24 hpi measured using ELISA. (E) Monocyte recruitment into VACV-infected ears of IFNαR^-/-^ mice 5 dpi assayed by flow cytometry, n=4 mice per group. (F) Ly6G^+^ cell recruitment into VACV-infected ears of IFNαR^-/-^ mice 5 dpi measured by flow cytometry, n=4 mice per group. (G) Immunofluorescence microscopy of IFNαR^-/-^ ears 5 dpi with VACV-CCL4 or VACV-mCherry for monocyte localization (Ly6C-Blue) in relation to VACV-infected cells (mKate/mCherry-Red). (H-J) Tissue pathology of the ear pinnae of IFNαR^-/-^ mice after intradermal infection with VACV; ear thickness (H), lesion size (I), or total tissue damage (J), n=6-7 mice per group. (K) Representative pictures of ear pinnae of IFNαR^-/-^ mice following intradermal VACV infection.

## Discussion

The importance of T1-IFN in control of virus replication has led to the evolution of a myriad of evasion strategies employed by viruses across multiple families, aimed at allowing virus replication and propagation to a new host. However, using a virus system in which multiple stages of T1-IFN induction and downstream signaling are inhibited, our data clearly indicate that, despite a lack of induction of T1-IFN within infected cells *in vitro*, T1-IFN is induced and acts to moderate tissue pathology following virus infection *in vivo*. Our data support a role for multiple host mechanisms that allow control of virus-dependent pathology but are independent of control over virus replication. Importantly, these mechanisms occur in the face of multiple, effective evasion strategies deployed by the viral pathogen.

Induction of T1-IFN following VACV infection *in vivo* occurred via the cytosolic DNA sensor, cGAS and its adapter protein STING. VACV completely blocks cGAS and STING-dependent induction of T1-IFN using multiple mechanisms *in vitro*, including expression of B2, F17, C4, C16 and C6 [15, 16, 17 Eaglesham, 2019 #43050]. However, production of both T1-IFN mRNA and protein, as well as protection against enhanced tissue pathology following VACV infection, was dependent upon cGAS, but not upon other PRR signaling pathways. This contrasts with other studies, which have demonstrated that cGAS is required to protect against VACV challenge, but which have stopped short of demonstrating a role for cGAS in T1-IFN production following VACV infection [18, 37, 38]. The lack of a link between cGAS and T1-IFN production following VACV infection *in vivo* likely stems from technical issues, and we developed a number of highly sensitive *ex vivo* approaches in order to be able to detect T1-IFN transcripts and protein from VACV-infected tissue. Using these approaches, we clearly demonstrate that cGAS signaling is required for T1-IFN induction following VACV infection. We found that the majority of infected cells *in vivo* mirror our *in vitro* finding, and do not express T1-IFN mRNA, likely due to the action of multiple VACV immunomodulators. We anticipated that the induction of T1-IFN that we observed *in vivo* might be the result of the transfer of cGAS-produced cGAMP to neighboring uninfected cells [60, 61]. Once within neighboring cells, cGAMP could induce T1-IFN production via STING within cells that are not subject to immunomodulatory VACV proteins, allowing for efficient expression. However, all of the IFN-β mRNA we detected was present within a small population of VACV-infected cells that appeared functionally distinct from the majority of infected cells. Therefore, a specialized population of cells may allow the host to subvert viral evasion mechanisms and express protective T1-IFN. Therapeutic targeting of such a population during infection or vaccination may preferentially allow for the generation of a strong T1-IFN response within a tissue.

What would be the characteristic of an infected cell that can subvert the action of the viral B2, F17 and C6 proteins to allow for cGAS-dependent T1-IFN expression? T1-IFN expression begins by day 3 post-infection, a time point when inflammatory monocytes are the first recruited cells to accumulate at the site of VACV infection. However, induction of T1-IFN still occurred in the absence of inflammatory monocyte recruitment (not shown). This was not unexpected, as T1-IFN was required for effective inflammatory monocytes recruitment. We have previously described T1-IFN production by the tissue protective Ly6G^+^ population of recruited cells [50]. However, the Ly6G^+^ population does not begin to accumulate until day 5 post-infection and does not peak until day 8-10 post-infection [50] i.e. significantly after the peak of T1-IFN production. Therefore, Ly6G^+^ cells may contribute to T1-IFN production after the peak of VACV replication, but they do not play a role in CCL4-mediated recruitment of classical inflammatory monocytes, which precede them into the VACV lesion. Our data are indicative of a role for tissue-resident VACV-infected cells in cGAS-mediated T1-IFN production. These cells could be specialized populations of keratinocytes, dermal fibroblasts, Langerhans cells, dermal DCs, tissue resident macrophages, γδ or other tissue-resident T cells, ILCs or even mast cells. Identification of the cells responsible for T1-IFN production following VACV infection will require extensive further studies.

The mechanism by which infected cells can produce T1-IFN in a cGAS-dependent manner *in vivo* in the face of viral evasion strategies that efficiently block this pathway *in vitro* is particularly interesting. The most effective way to bypass the blockade of cytosolic sensing within infected cells would be to inhibit expression of the viral proteins that target the cytosolic sensors. However, on a global level, the breadth of mechanisms employed by viruses to block cytosolic sensing likely precludes this approach. Indeed, in the system examined here, both the cGAS/STING and MDA5/MAVS pathways are targeted by multiple VACV proteins including B2, F17, K13, C6, E3 and K3 [15-17, 62-64]. Therefore, it is more likely that multiple cGAS/STING sensing and signaling pathways exist in specialized cells. In the case of the RNA-sensing pathway, MAVS signaling can occur from multiple sites within an infected cell, namely from the mitochondria or peroxisomal membranes [65]. These differential signaling sites produce different functional outcomes [65], raising the possibility that they may be differentially sensitive to viral inhibition. This would be the case if viral proteins are targeted to the mitochondria or ER, respectively, to inhibit the STING and MAVS signaling pathways. The possibility that these pathways may be differentially expressed, or perhaps enhanced, within specific cell populations is intriguing. Targeting of these specialized pathways may preferentially allow for the generation of a strong T1-IFN response during vaccination or therapeutic treatments.

The majority of T1-IFNs produced during VACV infection were IFN-β and IFN-α4, each of which are produced within the first round of T1-IFN induction, without the need to first induce expression of further ISGs such as *irf7*. Localization of IFN-β exclusively within VACV-infected cells implies that nearby cells are not triggered to produce either IFN-β or IFN-αs that are downstream from IFNαR in the positive feedback loop of T1-IFN induction. This observation supports two conclusions. First, the soluble T1-IFN binding protein encoded by VACV, B18 [66], appears to be very effective at preventing the spread of T1-IFN induction to nearby cells. Second, the majority of both IFN-β and IFN-α4 induction within infected cells is dependent upon IFNαR, STAT1 and STAT2, despite the inhibitory effect of the viral VH1 protein upon STAT1 activation [67]. These findings indicate that some level of positive feedback signaling through IFNαR in VACV-infected cells is required. B18 binds to the surface of uninfected cells and soaks up excess T1-IFN, preventing activation of these cells through IFNαR [66]. Therefore, B18 may not effectively bind to the surface of infected cells, perhaps due to virus-induced changes in the cell surface architecture and proteome.

It was surprising that VACV replication was unaffected by T1-IFN signaling. However, upregulation of antiviral ISGs was independent of T1-IFN in the infected ear, despite the observation that T1-IFN expression occurred only within infected cells. Many of the ISGs induced by T1-IFN signaling are important mediators in restricting the ability of viruses to infect and replicate within cells, although many have not been studied in the context of a VACV infection. However, during infection with VACV ISG15, which is strongly induced in a T1-IFN-independent manner in infected skin, controls mitochondrial function and limits virus replication within infected cells [68, 69]. The only logical explanation from these opposing observations is that ISG expression is induced by redundant signaling pathways via the expression of other cytokines. It is possible that other families of interferon species may induce the expression of ISGs in VACV-infected skin. In addition, there is significant crosstalk between the NF-κB and T1-IFN signaling pathways [44], so it is possible that other pathways are activated to stimulate ISG induction. NF-κB signaling is also inhibited in VACV-infected cells [28, 70, 71], but our profiling of the cytokine and chemokine response in the infected ear demonstrates strong upregulation of a number of inflammatory cytokines. We were unable to examine the induction of cytokines, chemokines or ISGs within infected vs uninfected cells *in situ*, but it is likely that upregulation of these effector molecules occurs within uninfected cell populations. Irrespective of the mechanisms by which ISGs are induced, it is nonetheless clear that antiviral ISGs do not play a role in the protection of infected tissue by T1-IFN following VACV infection.

Here, the action of T1-IFN is mediated through the CCL4-dependent recruitment of inflammatory monocytes, rather than through the direct inhibition of virus replication via the action of intracellular ISGs, a process that can be subverted by viral proteins within infected cells. In a number of viral infections, including the VACV system used here, tissue-resident macrophage populations in secondary lymphoid organs are required to prevent systemic dissemination of viruses [53, 56-58, 72]. After infection with some viruses, infection of macrophage populations is required in order to facilitate development of an effective adaptive immune response [55]. However, the role of recruited inflammatory monocyte populations after infection differs from that of macrophages resident in systemic organs [53]. Although inflammatory monocyte recruitment has been indirectly implicated in the response to virus infection [73-75], the mechanisms employed by recruited inflammatory monocyte populations are unknown. Monocytes do not directly inhibit VACV replication within other infected cell populations, as there is no large change in VACV titer, despite the marked change in tissue pathology in the absence of T1-IFN signaling. Infectious VACV can be recovered from inflammatory monocytes themselves when they are sorted from VACV-infected ears [19]. However, the titer recovered from inflammatory monocytes was less than 2 pfu per cell, and could represent input levels of virus rather than progeny virions. Here, we show that monocyte recruitment to the VACV-infected ear was dependent upon T1-IFN mediated induction of CCL4, although the requirement was incomplete. However, in the absence of CCL4 induction, recruitment to the VACV lesion was essentially completely ablated. As the majority of inflammatory monocytes recruited to the VACV lesion become infected, it is likely that infection of these cells may play a role in the tissue-protective phenotype that they mediate. Here, we demonstrate that inflammatory monocytes are infected, but the blockade in DNA replication in the majority of cells prevents the production of infectious virions. Defective recruitment of monocytes may account for the slight increase in VACV titer we recovered from cGAS^-/-^ and IFNαR^-/-^ mice compared to wild-type mice, as we observed a similar increase in titer in mice in which we have used multiple methodologies to deplete monocytes at the site of VACV infection, but which retain intact T1-IFN induction [53]. This further suggests that a lack of T1-IFN does not render ear tissue more permissive to VACV replication. We propose that inflammatory monocytes at the site of infection can function as a virus sponge, thereby limiting the proportion of ear tissue that is infected, and subsequently eliminated by the adaptive immune response. Indeed, surrounding ear tissue from IFNαR^-/-^ mice had a greater proportion of VACV-infected cells compared to wild-type mice, but VACV titers were not substantially increased, supporting this model. It is also possible that inflammatory monocytes may facilitate a response to “wall-off” the infected tissue and promote local wound healing responses. In either way, inflammatory monocytes can moderate virus-dependent tissue pathology in a T1-IFN-dependent manner that is independent of the induction of classical antiviral ISGs or inhibition of virus replication.

Taken as a whole, our findings provide a more complete picture of the way in which the host uses multiple mechanisms to bypass a myriad of viral evasion strategies, and thus still produce and utilize T1-IFN in a protective antiviral capacity. Induction of T1-IFN requires infection of specialized populations of tissue-resident cells that can subvert the viral blockade of cGAS/STING sensing. The infected cell population only produces “early” species of T1-IFN, likely due to the effects of soluble T1-IFN binding proteins and a blockade in STAT1 signaling within infected cells. However, the small amount of T1-IFN made is sufficient to induce a positive feedback loop of T1-IFN within infected cells to stimulate downstream expression of *ccl4* and subsequent recruitment of inflammatory monocytes. The inflammatory monocytes become the major infected cell population within the lesion, but are non-productively infected, effectively throwing themselves on the exploding grenade of the infection.

## Materials and Methods

### Mice

Mice were on the C57BL/6 background unless otherwise stated. Wild-type (catalog no. 00664), cGAS^-/-^ (catalog no. 026554), STING^gt/gt^ (catalog no. 017537), MyD88^-/-^ (catalog no. 009088), TRIF^Lps/Lps^ (catalog no. 005037), MAVS^-/-^ (catalog no. 008634 on the B6.129 background), STAT2^-/-^ (catalog no. 023309), and STAT1^+/-^ (catalog no. 012606, subsequently bred to produce homozygous STAT1^-/-^) mice were purchased from the Jackson Laboratory and housed at the Penn State College of Medicine. IFNαR^-/-^ (Jax catalog no. 32045) mice were a gift from Dr. Ziaur Rahman (Penn State College of Medicine). All mice were maintained in the specific-pathogen-free facility at the Penn State College of Medicine. *In vivo* experiments utilized 6-12-week-old mice. All animals were treated in accordance with the National Institutes of Health and AAALAC International Regulations. The Penn State College of Medicine Institutional Animal Care and Use Committee approved all experiments.

### Ethics Statement

All animals were maintained in the specific-pathogen-free facility of the Hershey Medical Center and treated in accordance with the National Institutes of Health and AAALAC International regulations. All animal experiments and procedures were approved in protocols 47731, 47033, 47597, 46111 and 44723 by the Penn State Hershey Institutional Animal Care and Use Committee (Animal Welfare Assurance # A3045-01) that follows the Office of Laboratory Animal Welfare PHS Policy on Humane Care and Use of Laboratory Animals, 2015.

### Viruses and cell lines

VACV (strain Western Reserve), or recombinant viruses (see below), were grown and purified as previously described [53]. Heat-inactivation of VACV was achieved by incubating aliquots of virus at 56°C in a shaking incubator for 1 hour just prior to infection. To easily identify VACV-infected cells *in vivo*, mice were infected with a recombinant VACV expressing influenza nucleoprotein tagged with eGFP, in which fluorescent protein accumulates in the nucleus of infected cells (VACV-NP-S-eGFP) [72]. In order to identify virus-encoded RNA, we used VACV that expresses the ovalbumin gene from *Gallus gallus* (VACV-OVA) [76]. VSV was a kind gift from the late Dr. Leo Lefrancois and was grown and titered on BHK cells using standard protocols.

To identify infected cells in which abortive or productive infection was proceeding, we utilized VACV-eGFP-OVA that expresses eGFP coupled with OVA under the control of either the early/late p7.5 promoter (VACV Early-GFP) or the late promoter p11 (VACV Late-GFP). VACV-eGFP-OVA early and late viruses were prepared as follows. The plasmid pRB21 expressing the full-length vp37 VACV ORF with the p7.5 early/late promoter was a kind gift from Dr. Bernard Moss (Laboratory of Viral Diseases, MIAID, Bethesda, MD). The peGFP-C1 plasmid expressing OVA was a kind gift from Dr. Kenneth Rock (Department of Pathology, University of Massachusetts Medical School, Worcester, MA). pRB21 DNA was ligated with eGFP-OVA using T4 Ligase (Invitrogen). To make rVACV-eGFP-OVA with a late promoter, the p11 promoter was inserted in place of p7.5. All plasmid DNA was sequenced to ensure that the promoter, and eGFP-OVA sequences were correct. rVACV-eGFP-OVA-Early and rVACV-eGFP-OVA-Late were generated by transfecting plasmid DNA into BSC-1 cells infected with VACV-vRB21 at an MOI of 1 using the CellPhect Transfection Kit (GE Healthcare). The resulting rVACVs were plaque-purified 3× prior to characterization. The resulting rVACV-eGFP-OVA-Early and rVACV-eGFP-OVA-Late produced green fluorescence upon infection of WT3 cells and sequencing revealed the presence of the correct promoter and OVA sequences in DNA purified from virions.

To make a virus that expresses CCL4, a construct was ordered that contained 357 nucleotides of the 3’ end of the J4 open reading frame (ORF) followed by the early VACV promoter C6, the coding sequence for mKate2 and mouse CCL4 ORFs connected by an in-frame sequence for the P2A peptide. The CCL4 ORF was followed by a 3x VACV early transcription stop signal and 400 nucleotides of the 3’ end of the J5 ORF (IDT). These constructs were amplified by standard PCR and cloned into pCR2.1 and sequenced to ensure fidelity. Recombinant viruses were generated using homologous recombination by infecting HeLa cells with VACV WR strain and transfecting in the constructs (VACV-CCL4). Recombinant viruses were isolated based on their ability to produce red fluorescent plaques. The isolated recombinants were plaque-purified 4x before being amplified to produce working stocks. The integrity of the inserts was verified by sequencing. For microscopy experiments utilizing VACV-CCL4, we used VACV mCherry Ub-SIINFEKL (VACV-mCherry) [77] as the control for expression of a red fluorescent protein without CCL4.

The murine keratinocyte cell line 308 was originally derived from BALB/c mouse skin initiated with DMBA [78]. NR9456 is a murine macrophage cell line derived from differentiated bone marrow-derived cells immortalized by retroviral transformation [79] and was obtained through BEI Resources, NIAID, NIH. 143B cells are a thymidine kinase-deficient human osteosarcoma cell line. 143B and NR9456 cells were grown at 37°C in DMEM supplemented with 10% FBS, 20mM HEPES, 1.5 g/L NaHCO_3_, 0.3 g/L L-glutamine, and 0.1 g/L penicillin and streptomycin. 308 cells were grown at 37°C in DMEM containing 4.5 g/L glucose, L-glutamine and sodium pyruvate supplemented with 10% FBS, 0.1 g/L penicillin and streptomycin and an additional 0.3 g/L L-glutamine.

### VACV infection

For intradermal VACV infection, 10^4^ pfu VACV in a 10μl volume of Hank’s Balanced Salt Solution (HBSS)/0.1% bovine serum albumin (BSA) was injected into the middle of each ear pinnae [21]. Intradermal infection causes substantial tissue loss at late times post-infection, preventing accurate measurement of titers at those times. Therefore, to assess virus titer at late times post-infection, mice were infected by scarification of the ear pinnae. Briefly, a drop of HBSS/0.1%BSA containing 10^4^ pfu VACV was placed on the center of the ear pinnae and a 28-gauge needle was used to scratch the ear through the droplet [80]. To produce a localized infection suitable for microscopy imaging, mice were infected by dipping a bifurcated needle into HBSS/0.1%BSA containing 2×10^8^ pfu/ml VACV and then gently puncturing each ear pinnae ten times [19]. To evaluate tissue pathology during VACV infection, ear thickness was measured daily using a 0.0001m dial micrometer (Mitutoyo). After the appearance of a visible lesion, the diameter of the lesion or of areas of tissue loss were measured [50].

For infection of cells *in vitro*, 5e^5^-1e^6^ cells were seeded onto 24-well plates 24 hours prior to infection. Cells were infected at an MOI of 10 in 250μl HBSS/0.1%BSA for 1 hour at 37°C, inoculum was removed and replaced with 500μl of pre-warmed media and placed at 37°C for either an additional 5 or 23 hours.

### VACV titer

Virus replication was quantified by plaque assay as previously described [50, 53]. Infected ears were harvested at the times indicated and homogenized using a TissueLyser II (Qiagen), then centrifuged for 2 minutes at 12,000 × g. Supernatant was transferred into a fresh tube and virus titer was determined by plaque assay on 143B cells.

### RNA isolation and RT-qPCR

For gene expression *in vitro*, cells were infected as described above. After 6 hours total infection time, media was removed and cells were lysed with 500μl TRIzol reagent (Invitrogen), transferred to a 1.7ml tube, and placed at −20°C until RNA extraction. For RNA isolation, 100μl chloroform was added to thawed tubes, briefly vortexed and centrifuged at 12,000 × g for 15 minutes at 4°C. The upper aqueous phase was transferred into a clean tube, 1 volume of 70% EtOH was added, and RNA was purified using the RNeasy mini kit (Qiagen) with on-column DNase-I treatment.

For gene expression *in vivo*, ears were harvested from mice at the times indicated post-infection, flash-frozen in liquid NO_2_ and placed at −80°C. For RNA isolation, ears were homogenized using a TissueLyser II (Qiagen) and RNA was purified from tissue lysate using the RNeasy Mini Kit (Qiagen) treated with DNase-I (Qiagen) in-solution.

For qPCR using Taqman Gene Expression Assays (Applied Biosystems) or Universal Probe Library (Roche), cDNA was prepared using the Hi-Capacity cDNA Synthesis Kit (Applied Biosystems). For qPCR using RT^2^ PCR Profiler Arrays (Qiagen), cDNA was prepared using RT^2^ First Strand Kit (Qiagen). qPCR was carried out on a StepOnePlus (Applied Biosystems) with either FastStart Universal Probe Master Mix (Roche) or RT^2^ SYBR Green qPCR Master Mix (Qiagen). For Taqman and FastStart Universal Probe assays, changes in gene expression are expressed as fold change using the ΔΔ^Ct^ calculation method against naïve mice of the same genotype or mock-infected cells with *gapdh* as the housekeeping gene. For RT^2^ PCR Profiler Array data, changes in gene expression are displayed as mean fold change between groups of mice relative to a panel of “housekeeping” genes. Amplification of bulk IFN-α transcripts was detected using Universal Probe Library probe #76 (Roche) with forward primer ARSYtgtStgatgcaRcaggt and reverse primer ggWacacagtgatcctgtgg; and was compared to amplification of *gapdh* using Universal Probe Library probe #80 (Roche) with forward primer tgtccgtcgtggatctgac and reverse primer cctgcttcaccaccttcttg. Primers were from Integrated DNA Technologies.

### ELISA

For detection of protein from cells infected *in vitro*, cells were infected as described above. After the initial 1 hour of infection, the inoculum was removed and replaced with 500μl of pre-warmed media supplemented with 2% FBS and cells were incubated at 37°C for an additional 23 hours. After 24 hours of total infection time, media was transferred into tubes containing phenylmethylsulfonyl fluoride (PMSF) (Sigma) and a Protease Inhibitor Cocktail (Sigma) and placed at −80°C until the ELISA was performed.

For detection of protein from VACV-infected ear tissue, ears were infected and harvested as described above. Frozen ears were homogenized in PBS containing PMSF (Sigma) and Protease Inhibitor Cocktail (Sigma) using a TissueLyser II (Qiagen), centrifuged for 3 min at 12,000 × g and the lysate was transferred to a clean tube.

IFN-β protein was detected using the mouse Hi-Sensitivity IFN-β ELISA kit (PBL Assay Science). CCL4 protein was detected using the mouse CCL4/MIP-1 beta Quantikine ELISA kit (R&D Systems). ELISAs were carried out according to manufacturer’s specifications.

### Flow cytometry

For the detection of the mKate2 transgene, 2e^6^ 143B cells were infected in 5ml tubes at an MOI of 10 in 500μl HBSS/0.1%BSA, incubated in a 37°C water bath for 1 hour, 3 ml of pre-warmed media was added, and tubes were incubated in a rotating incubator at 37°C for an additional 5 hours. Cells were washed and samples were acquired on a LSRII flow cytometer (BD Biosciences), data were analyzed using FlowJo software (TreeStar).

Ears were harvested and processed as previously described [50, 53]. Briefly, ears were separated into dorsal and ventral halves and digested in 1 mg/ml collagenase XI (Sigma) for 60 minutes at 37°C and a single cell suspension made. Live cells were incubated in Fc block and 10% normal mouse serum (Gemini Bio-Products) at 4°C, and then stained in that solution. Cells were stained with antibodies recognizing Ly6C (clone HK1.4), CD11b (clone M1/70), Ly6G (clone 1A8), CD45 (clone 30-F11), CD64 (clone X54-5/7.1), CD11c (clone N418), CCR5 (clone HM-CCR5), CXCR3 (clone CXCR3-173), CXCR5 (clone L138D7), Streptavidin from BioLegend, CD19 (clone 1D3), CD90.2 (clone 53-2.1), and NK1.1 (clone PK136) from BD Biosciences, and CCR2 (clone REA538) from Miltenyi Biotec. Samples were acquired on a LSRII flow cytometer (BD Biosciences), and data were analyzed using FlowJo software (TreeStar). Inflammatory monocytes were gated as CD45^+^ (CD19, CD90.2, NK1.1)^-^ CD11b^+^ Ly6C^hi^ Ly6G^-^ CD11c^-^ CD64^+^ [53]. Non-classical tissue-protective monocytes were gated as CD45^+^ (CD19, CD90.2, NK1.1)^-^ CD11b^hi^ Ly6C^+^ Ly6G^+^ [50].

### Fluorescent imaging

Mice were infected with VACV in the center of each ear pinnae using a bifurcated needle. Five days post-infection, ears were harvested and embedded in Tissue-Tek OCT (Sakura Finetek), rapidly frozen by immersion in liquid nitrogen-cooled 2-methyl butane, and stored at −80°C. Lateral cryostat sections (10 μm) were cut at −20°C, mounted on glass slides and fixed for 30 min in 4% paraformaldehyde in PBS (pH 7.4). For both assays, three-dimensional images were collected on an Olympus IX81 fluorescent microscope, deconvolved and analyzed using Slidebook 6.0 software (Intelligent Imaging Innovations).

For RNA fluorescence *in situ* hybridization, mice were infected with VACV-OVA. To visualize plasma membranes, tissue was stained with Wheat Germ Agglutinin conjugated to CF488 (1:1000, Biotium). Tissue was fixed again and RNA was visualized using the ViewRNA Cell Plus Assay (Invitrogen) with probes recognizing murine IFN-β and gallus ovalbumin (Affymetrix) according to manufacturer’s specifications with the omission of using a protease. Nuclei were stained with DAPI.

For Immunofluorescence microscopy, mice were infected with VACV-NP-S-eGFP, VACV-CCL4, or VACV-mCherry. Tissue was stained with antibodies to Ly6C (clone HK1.4 1:2500, Abcam) and visualized using secondary donkey anti-rat Alexa 647 (1:500, Jackson Immunoresearch) or donkey anti-rat BV421 (1:100, Jackson Immunoresearch).

### Data analysis

Data were analyzed and graphed using Excel (Microsoft), Prism (Graphpad), and RT^2^ Profiler Array web-based software (Qiagen). Data are expressed as mean ± standard error of the mean, with individual data points shown where applicable. RT^2^ PCR Profiler Array data are expressed as mean fold change vs. p-value. Means were compared using an unpaired, two-way student’s t-test or two-way ANOVA as applicable. Data represents at least three individual experiments. Significance between groups was determined by p-value <0.05, and is displayed as; * = p<0.05, ** = p<0.01, *** =p<0.001, **** = p<0.0001.

